# From epigenetic landscape to phenotypic fitness landscape: evolutionary effect of pathogens on host traits

**DOI:** 10.1101/051904

**Authors:** Mark Jayson V. Cortez, Jomar F. Rabajante, Jerrold M. Tubay, Ariel L. Babierra

## Abstract

The epigenetic landscape illustrates how cells differentiate into different types through the control of gene regulatory networks. Numerous studies have investigated epigenetic gene regulation but there are limited studies on how the epigenetic landscape and the presence of pathogens influence the evolution of host traits. Here we formulate a multistable decision-switch model involving many possible phenotypes with the antagonistic influence of parasitism. As expected, pathogens can drive dominant (common) phenotypes to become inferior, such as through negative frequency-dependent selection. Furthermore, novel predictions of our model show that parasitism can steer the dynamics of phenotype specification from multistable equilibrium convergence to oscillations. This oscillatory behavior could explain pathogen-mediated epimutations and excessive phenotypic plasticity. The Red Queen dynamics also occur in certain parameter space of the model, which demonstrates winnerless cyclic phenotype-switching in hosts and in pathogens. The results of our simulations elucidate how epigenetic landscape is associated with the phenotypic fitness landscape and how parasitism facilitates non-genetic phenotypic diversity.

## Introduction

The mechanisms of epigenetics are multifaceted and complex [Danchin et al. 2011; Gomez-Diaz et al. 2012; Huang 2013; Duncan et al. 2014; Kilvitis et al. 2014; Skinner 2015]. In spite of numerous experimental and theoretical studies on the epigenetic landscape of phenotype specification [Richards 2008; Rohlf et al. 2012; Huang 2013; Kilvitis et al. 2014; Rabajante et al. 2015], its behavior is not fully understood, especially in the presence of pathogens (e.g., parasites). Parasitism-mediated alterations in the dynamics of the epigenetic landscape have not gained high attention in computational systems biology [Poulin & Thomas 2008; Boyko & Kovalchuk 2011; Bierne et al. 2012; Gomez-Diaz 2012]. Particularly, mathematical models need to advance to capture the important qualitative behavior of the epigenetic landscape that involves many factors, such as diseases that affect phenotype specification and fitness of species [Raghavan et al. 2010; Geoghegan & Spencer 2011; Huang 2012; Rabajante & Babierra 2015; Rabajante et al. 2015]. Theoretical predictions that employ epigenetic perspective can guide ecological, agricultural, epidemiological and biomedical studies in investigating the implications of host-pathogen interaction in phenotype variation and trait heritability [Woolhouse et al. 2002; Mallard & Wilkie 2007; Poulin & Thomas 2008; Kasuga & Gijzen 2013; Rabajante et al. 2015]. It is plausible to incorporate epigenetics to the study of parasite-induced evolution since the epigenetic pattern (epigenotype) in gene expression is a non-genetic factor that could be passed-on from parents to offspring [Pal & Miklos 1999; Bond & Finnegan 2007; Bonduriansky & Day 2009; Petronis 2010; Danchin et al. 2011; Kilvitis et al. 2014; English et al. 2015].

### Epigenetic attractors

Mathematical models of epigenetic landscape describe cell-fate determination/specification as a process converging to cellular attractors (see individual epigenetic landscapes in Fig. 1) [Cinquin & Demongeot 2005; Huang 2012; Furusawa & Kaneko 2012; Huang 2013; Rabajante & Babierra 2015; Rabajante & Talaue 2015]. Stem cell differentiation is illustrated as a branching progression from totipotency to various cell lineages to different terminally specialized cell types [Furusawa & Kaneko 2012; Huang 2013; Rabajante & Babierra 2015]. Through the regulation of gene interaction, cells decide where to converge from the multiple attractors present in the epigenetic landscape. An attractor can be any fate of the cell, such as metastable, terminally specialized, cancer, quiescent, senescence and apoptotic states [Furusawa & Kaneko 2012; Huang 2013; Li & Wang 2014; Marco et al. 2014; Rabajante & Babierra 2015; Rabajante et al. 2015]. Metastable states could be totipotent, pluripotent, multipotent or progenitor. The attractors that characterize terminally specialized cells represent different phenotypic fates (e.g., muscle, skin, blood, neuron, bone, fat). Moreover, cells may not track a one-directional linear pathway in the epigenetic landscape, that is, the pathway can be multidirectional or circular [Enver et al. 2009; Furusawa & Kaneko 2012; Rabajante & Babierra 2015]. Cells are said to be plastic and can be driven to undergo dedifferentiation (cells regress to earlier state, e.g., from specialized cells back to pluripotent state) and transdifferentiation (cells transfer to other lineages, e.g., from mesenchymal to neural lineage) [Jopling et al. 2011; Xu et al. 2014]. Considerable degree of biological noise and induction by external stimuli may also affect the direction of the developing cells [Rabajante & Babierra 2015; Rabajante & Talaue2015].

**Fig. 1.**
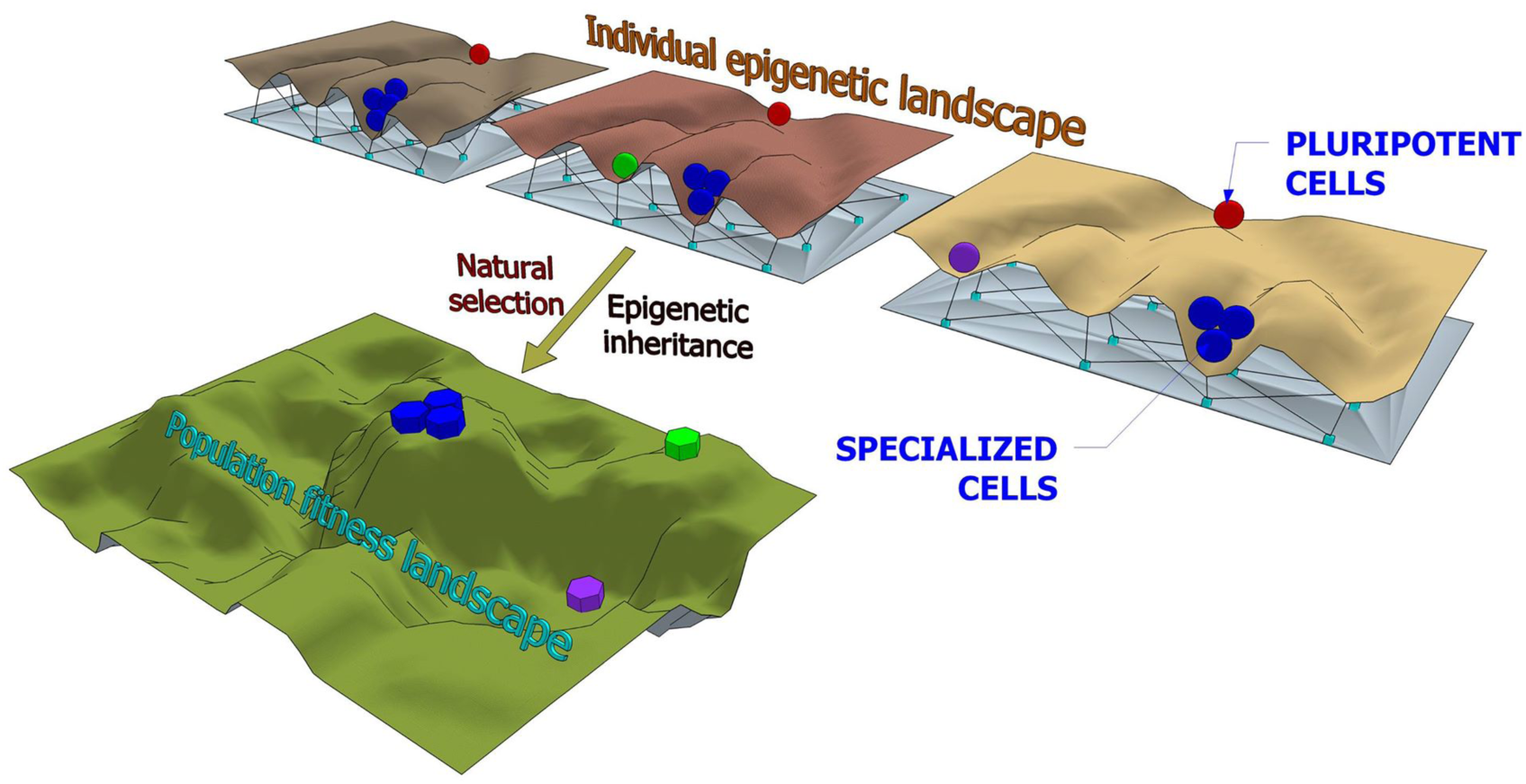
The link between the epigenetic landscape and fitness landscape. In our case, natural selection (e.g., negative frequency-dependent selection) is induced by environmental/external factors, such as parasitism [Poulin & Thomas 2008; Bonduriansky & Day 2009; Boyko & Kovalchuk 2011; Gomez-Diaz et al. 2012; Kasuga & Gijzen 2013; Soen 2014]. We assume that the collective dynamics of the individual epigenetic landscapes are reflected in the phenotypic fitness landscape at the population level through epigenetic inheritance (because of epigenetic memory) and parasitism-induced selection. If a phenotype is advantageous and there is adequate epigenetic memory, a population of species may select this phenotype resulting in higher phenotypic fitness.

The concept of cellular attractor is evolving and has various interpretations. However, a general rule for being an attractor is that the trajectory of developing cells in the epigenetic landscape from a neighborhood of initial conditions (basin of attraction) should converge to this attractor state [Huang 2012]. Any perturbation within the basin of attraction should not result in instability, that is, the state is resilient and homeostatic [Huang 2013]. Mathematically, the attractors are represented by stable equilibrium points [Huang 2013; Rabajante & Babierra 2015], stable limit cycles [Rabajante & Babierra 2015], strange chaotic attractors [Furusawa & Kaneko 2012] or noisy attractors [Huang 2009]. Each attractor has its own basin of attraction, which can be of different sizes. Moreover, the parameter values in the mathematical models are dictated by the genes and their interaction. Gene interaction is not static but rather dynamic [Rabajante & Babierra 2015]. Modification in the parameter values may or may not preserve the stability of an attractor. Sometimes the modifications result in bifurcation that transforms the topography of the epigenetic landscape to form a new attractor or obliterate an existing attractor [Huang 2013; Rabajante & Talaue 2015]. Dynamically changing a parameter value could lead cells to change fates by moving their trajectory from one basin of attraction to another basin of attraction [Rabajante & Talaue 2015].

### Host-parasite coevolution

The interaction and coevolution between hosts and parasites are widely investigated. In evolutionary biology, arms race competition and the Red Queen dynamics have been hypothesized to cause coadaptation/coevolution in hosts and pathogens [Avrani et al. 2012; Brockhurst et al. 2014; Raberg et al. 2014; Rabajante et al. 2015; Voje et al. 2015]. For the host to survive parasitism, it increases its defense traits; but for the pathogen that relies heavily on the host, it needs to counteract the host defense and increase its pathogenicity. This may result in a winnerless arms race competition [Brockhurst et al. 2014; Rabajante et al. 2015]. In several cases where there are multiple types of hosts, parasitism can decrease the abundance of the common host type permitting a rare type to become the new dominant (negative frequency-dependent selection). Common host types, especially monocultures, are more susceptible to the attack of pathogens. This phenomenon is akin to the killing the winner hypothesis [Avrani et al. 2012; Rabajante et al. 2015]. Host-pathogen interaction is one of the explanations for the extinction and diversity of species [Miura et al. 2006; Brockhurst et al. 2014].

If negative frequency-dependent selection persists, it could result in fluctuating Red Queen dynamics which is illustrated by cyclic phenotype-switching [Brockhurst et al. 2014; Rabajante et al. 2015]. The fluctuating Red Queen dynamics is a candidate model of recurrent punctuated equilibrium process that is driven by biotic interaction [Rabajante et al. 2015]. This recurrent punctuated equilibrium process has out-of-phase heteroclinic cycles of rapid negative frequency-dependent selection, followed by stasis, then negative frequency-dependent selection again. Simulations show that the Red Queen dynamics are robust against certain level of noise but the ordering of cyclic phenotype-switching could vary, resulting in a diversified possibilities of evolutionary patterns [Rabajante et al. 2015].

### Bridging the gap

Phenotypic variation is affected by inherited and non-inherited biological information [Danchin et al. 2011; Day & Bonduriansky 2011; English et al. 2015; Skinner 2015; Soubry 2015]. Inherited information, which is influenced by genetic and non-genetic (e.g., epigenetic) factors, is important in evolution [Danchin et al. 2011; Richards et al. 2012; Klironomos et al. 2013; Smith & Ritchie 2013; Nishikawa & Kinjo 2014; Santos et al. 2015]. That is, genotype-by-epigenotype-by-environment regulates transmission of phenotypes from one generation to another (inclusive heritability) [Danchin et al. 2011; Gervasi et al. 2015]. Here we aim to contribute in bridging a gap between the study of epigenetics and evolutionary biology using a mathematical model, specifically by linking the epigenetic landscape at the individual level to the phenotypic fitness landscape at the population level (Fig. 1). There are early discussions about the connection between the dynamics of the epigenetic landscape and fitness landscape, such as in the case of tumor progression [Huang 2013], but here we consider the effect of parasitism as a major driver of phenotypic evolution. In Figure 2, the genotype-by-epigenotype interaction is represented by a phenotype decision-switch network where parasitism acts as the influencing environmental factor. In our mathematical model (see Methods), we assume that phenotype specification is determined exclusively by epigenetic mechanisms where genotypes are stable, i.e., the structure of gene regulatory network is fixed. This assumption allows us to decouple the effect of genetic and epigenetic factors in the genotype-by-epigenotype interaction, and focus our observation on the effects of parasitism-modified epigenotypes on host traits.

**Fig. 2.**
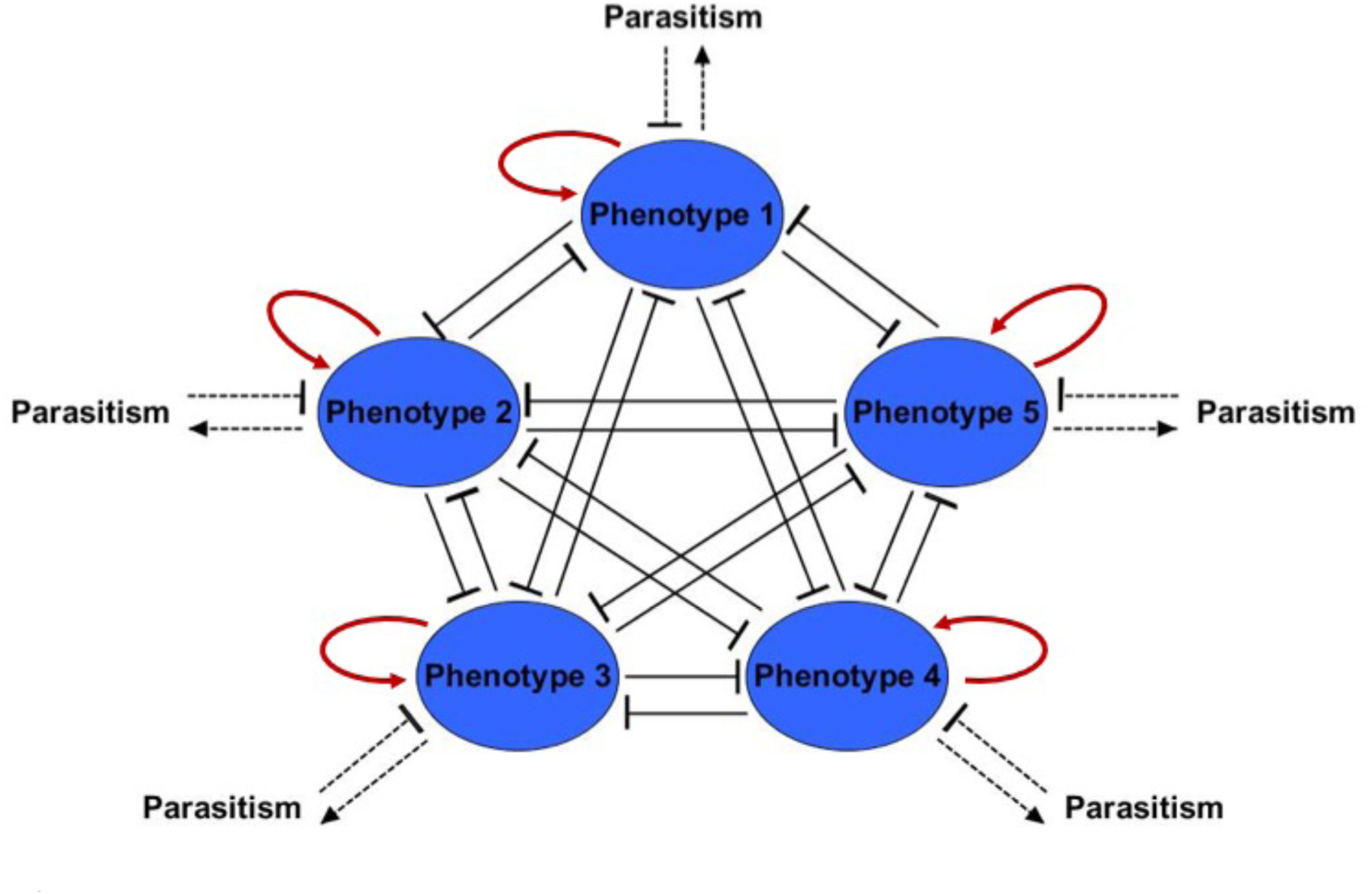
The phenotype decision-switch network. The qualitative dynamics of the gene regulatory network that determines the selected phenotypes follow this decision-switch network [Cinquin & Demongeot 2005; Macarthur et al. 2008; Rabajante & Babierra 2015]. Suppose there are 5 phenotypes (nodes). At the individual epigenetics level, the nodes have mutual repression because it is presumed that a specialized cell expresses a unique phenotype and inhibits the expression of the other phenotypes. The red positive feedback arrow represents self-stimulation. For simplicity, we assume that the structure of gene regulatory network is fixed, i.e., fixed parameter values (see early studies [Rabajante & Babierra 2015; Rabajante & Talaue 2015] for the simulation with dynamic gene regulatory network but without parasitism). Correspondingly, we assume that the average collective dynamics at the population level also adhere to such decision-switch network. Note that the frequency of phenotype is a continuum from 0 to 1. The frequency of a phenotype is negatively influenced by parasitism.

## Results and Discussion

At the cellular level, cells decide from multiple choices of phenotypes (epigenetic landscape in Fig. 1). In the presence of intense evolutionary pressures, such as presence of pathogens, genetic or non-genetic selection may occur [Danchin et al. 2011; Gomez-Diaz et al. 2012; Duncan et al. 2014; Kilvitis et al. 2014; Skinner 2015; Rabajante et al. 2015]. Our discussion focuses on non-genetic selection driven by epigenetic mechanisms with the influence of pathogens. If a phenotype is advantageous and there is adequate epigenetic memory [Pal & Miklos 1999; Bond & Finnegan 2007; Geoghegan & Spencer 2012; Gomez-Diaz et al. 2012; DUrso & Brickner 2014; English et al. 2015], a population of species may select this phenotype resulting in higher phenotypic fitness (Fig. 1). However, this selection process and a phenotype becoming prevalent within a population are not straightforward and strict conditions are necessary [Takahashi et al. 2010; Vermeij & Roopnarine 2013; Rabajante et al. 2015]. For example, parasitism resulting in negative frequency-dependent selection requires specific structure of functional response and combination of parameter values (e.g., Figs. 3 and 4) [Rabajante et al. 2015]. The characteristics of the hosts (e.g., growth rate and competitive ability) and pathogens (e.g., pathogenicity, specificity and death rate) dictate the phenotypic outcome.

**Fig. 3.**
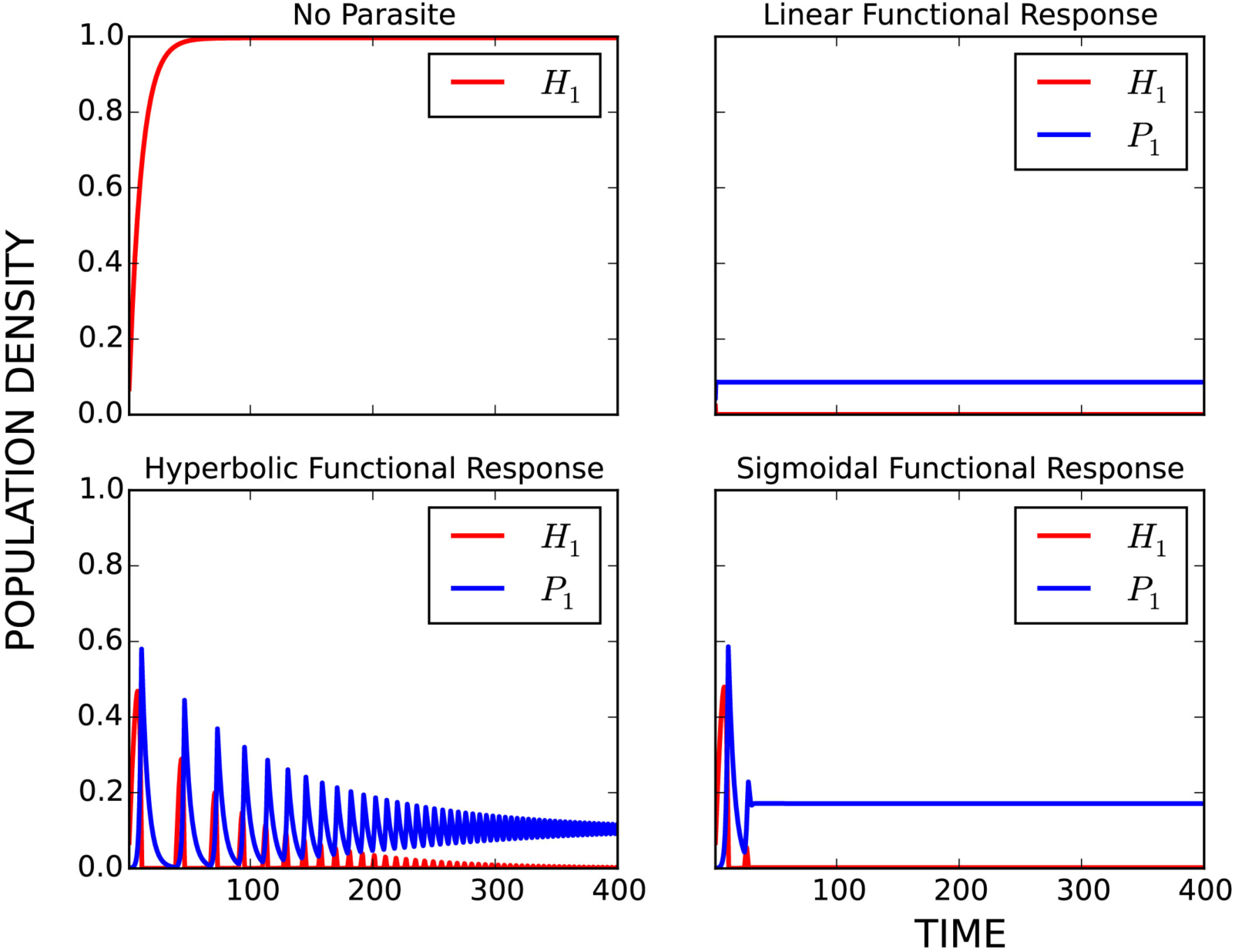
Effect of parasitism on host population. (**a**) Host population dynamics without parasitism. (**b-d**) Host population dynamics with parasitism using different functional response. Here host population density is defined as *Hi/K*, where *K* is the total carrying capacity. For comparison, we also assume that parasite population density is defined as *P_j_/K*.

Questions arise on whether it is the epigenetics structure that causes pathogenesis or whether the pathogens are the ones causing epigenetic changes [Gomez-Diaz et al. 2012]. We argue that these two questions are valid since the epigenome of hosts and the mechanisms of pathogens are components of an interacting system which is possibly influenced by feedback loops (Fig. 2). Particular parameters of the epigenetic model (Eq. 1 in Methods) may or may not allow disease to affect the outcome of gene regulation. On another side, the parameters (e.g., pathogenicity and specificity) and functional response in the parasitism model (Eq. 2 in Methods) dictate the severity of disease that can alter the topography of the epigenetic landscape.

### Equilibrium dynamics

As expected in many cases, coexistence of all phenotypes (sometimes with equal frequencies) and competitive exclusion occur (Fig. 4a and 4b). Coexistence represented by equilibrium point with all components having equal values may have limited range of initial value and parameter space, that is, some of the phenotypes could become inferior when conditions are changed. In competitive exclusion, there are dominant phenotypes and rare or extinct phenotypes in the long run. Competitive exclusion illustrates silencing or knockdown of some genes, and activation or sometimes overexpression of the other genes. In some cases, hosts are not able to regain growth, resulting in total silencing/extinction (Fig. 4c). The coexistence, competitive exclusion and total extinction of phenotypes are expected in a wide range of parameter space because the fundamental epigenetic model (Eq. 1 in Methods) describes a multistable system [Mostowy et al. 2012; Rabajante & Talaue2015].

**Fig. 4a.**
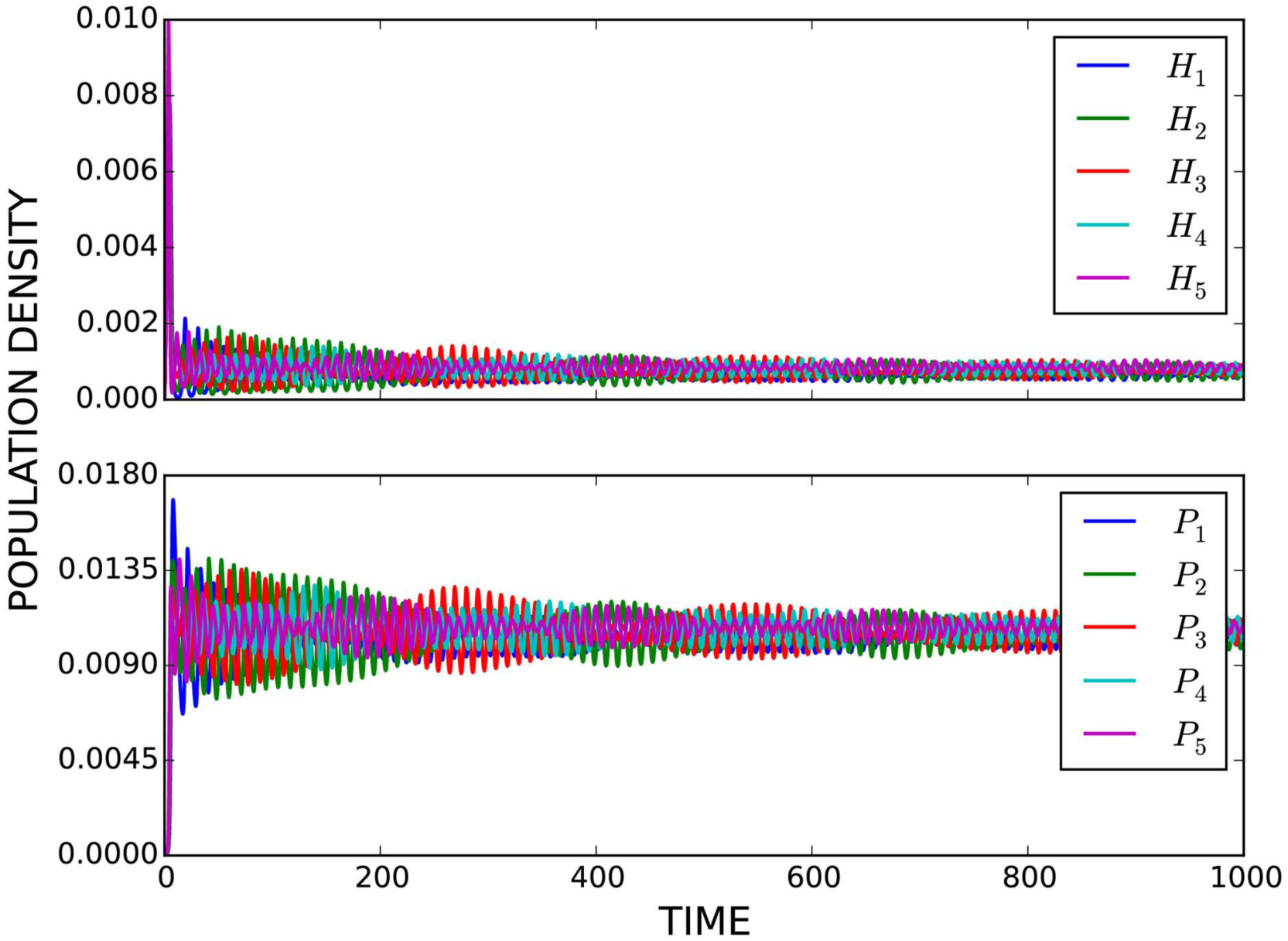
Different parasitism-induced population dynamics. Here host population density is defined as *H_i_/K*, where *K* is the total carrying capacity. For comparison, we also assume that parasite population density is defined as *P_j_/K*(**a**) Equilibrium-convergence showing coexistence of all host and parasite types.

**Fig. 4b.**
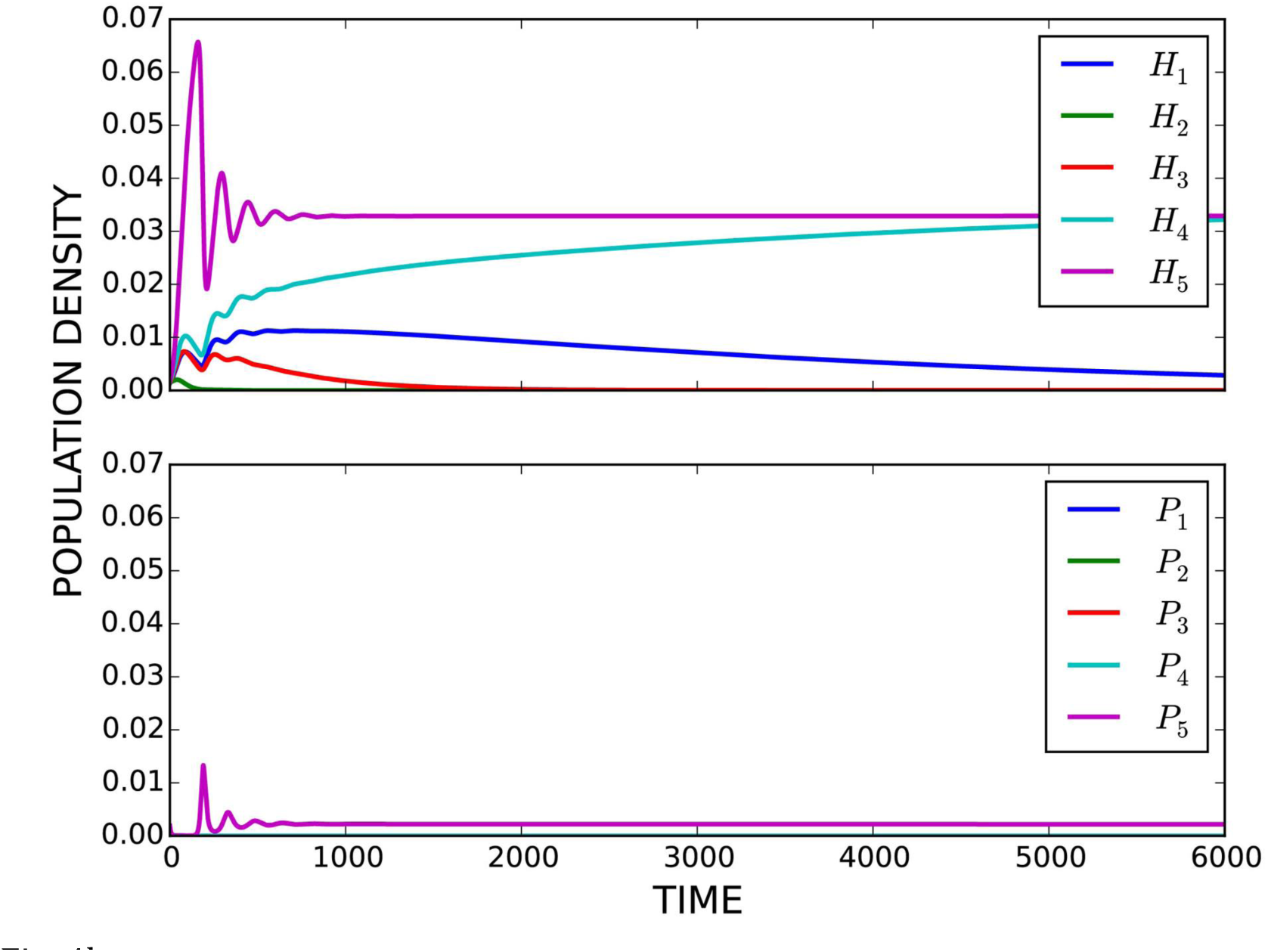
(**b**) Equilibrium-convergence showing competitive exclusion.

**Fig. 4c.**
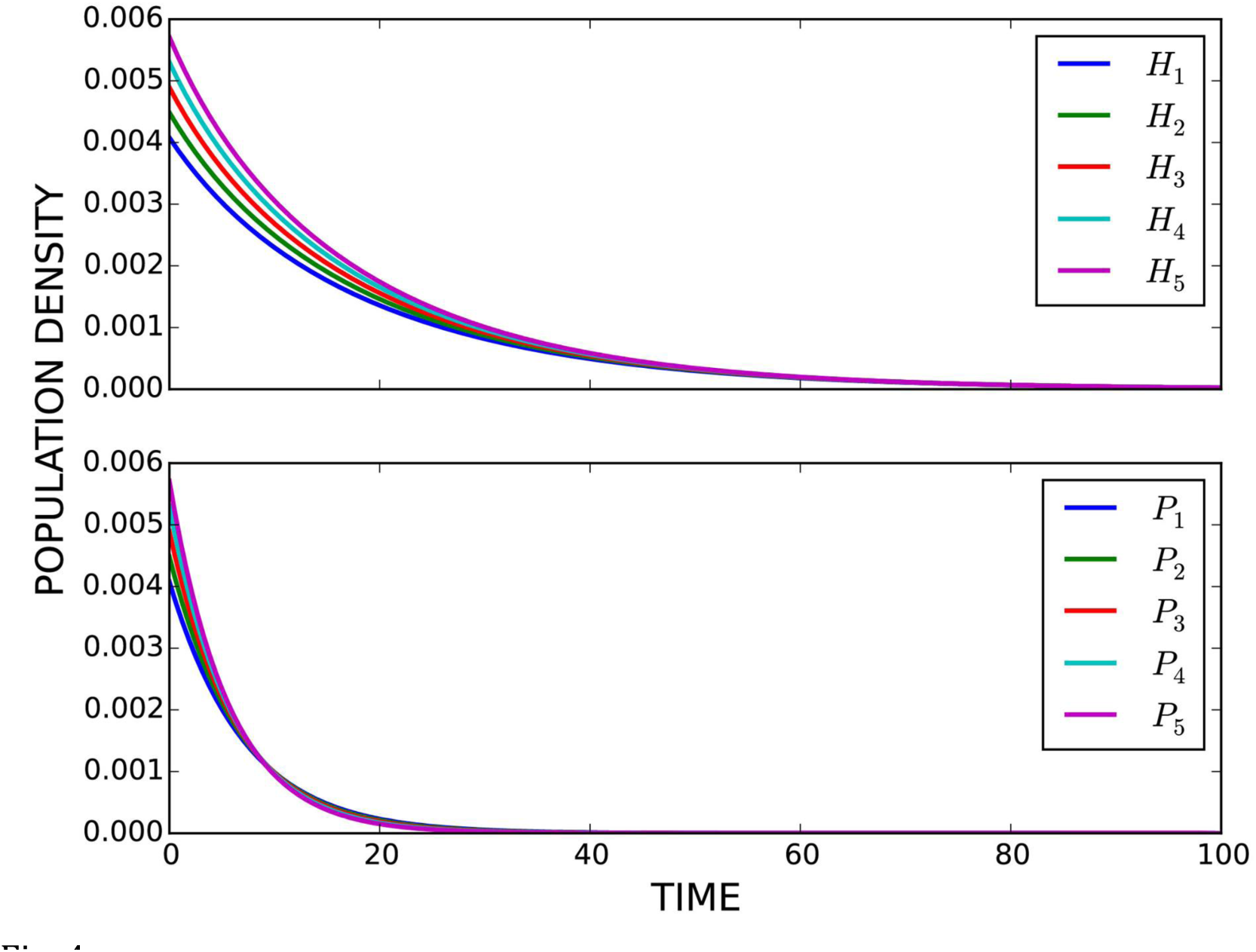
(**c**) Extinction of all host and parasite types.

Multistability pertains to the existence of many stable equilibrium points. Here multistability implies non-genetic variation. The meaning of non-genetic variation driven by epigenetic mechanisms is that two cells have identical DNA sequences (same set of genes) but they differ on what genes are expressed (e.g., activated or silenced during transcription) resulting in different phenotypes. For example, skin cells and muscle cells in an organism have similar set of genes because they came from one mother cell but they have different expressed genes that define the proteins specific for skin cells and specific for muscle cells. This idea can be extended to population epigenetics where non-genetic variation implies phenotypic heterogeneity not explained by genetics [Schlichting & Smith 2002; Johnson & Tricker 2010; Hughes 2012; Duncan et al. 2014; Kilvitis et al. 2014]. Non-genetic phenotypic heterogeneity is one of the survival strategies of species against environmental stress [Donaldson-Matasci et al. 2008; Soen 2014]. For example, some introduced species with low genetic variation can still become invasive because they can have high phenotypic diversity due to epigenetic variation [Kilvitis et al.2014].

### Oscillatory dynamics

Oscillations in phenotype frequencies are also possible to arise and they may characterize non-equilibrium polymorphism [Takahashi et al. 2010; Mostowy et al. 2012] and non-equilibrium phenotypic diversity (Figs. 4d and 4e). In fact, oscillatory behavior is a characteristic of stem cells and progenitor cells, which implies that oscillations have high potential to generate different cell types [Furusawa & Kaneko 2012; Rabajante & Babierra 2015]. In population biology point-of-view, if the minimum value of the oscillations are far from the edge of extinction, then oscillations exhibit permanent coexistence [Huisman & Weissing 2001]. If the minimum value of the oscillations are near the edge of extinction, then oscillations (Fig. 4e; e.g., the Red Queen dynamics) may lead to impermanent coexistence [Huisman & Weissing 2001]. It is called impermanent coexistence because random perturbations could drive the population frequency of a species to zero (guaranteed extinction); however, there are cases where Red Queen dynamics are robust against random noise [Rabajante et al. 2015].

**Fig. 4d.**
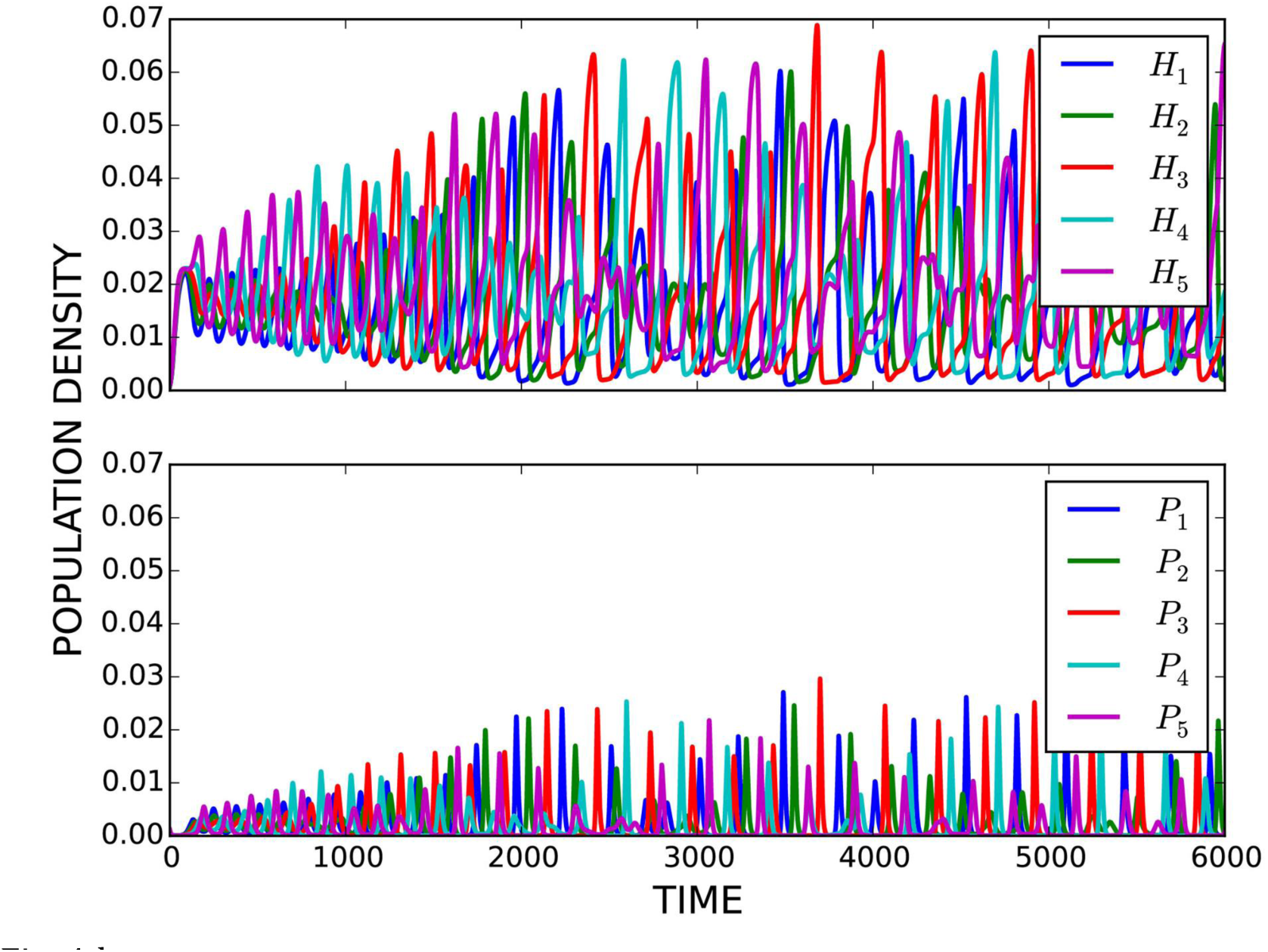
(**d**) Parasitism-induced oscillations.

**Fig. 4e.**
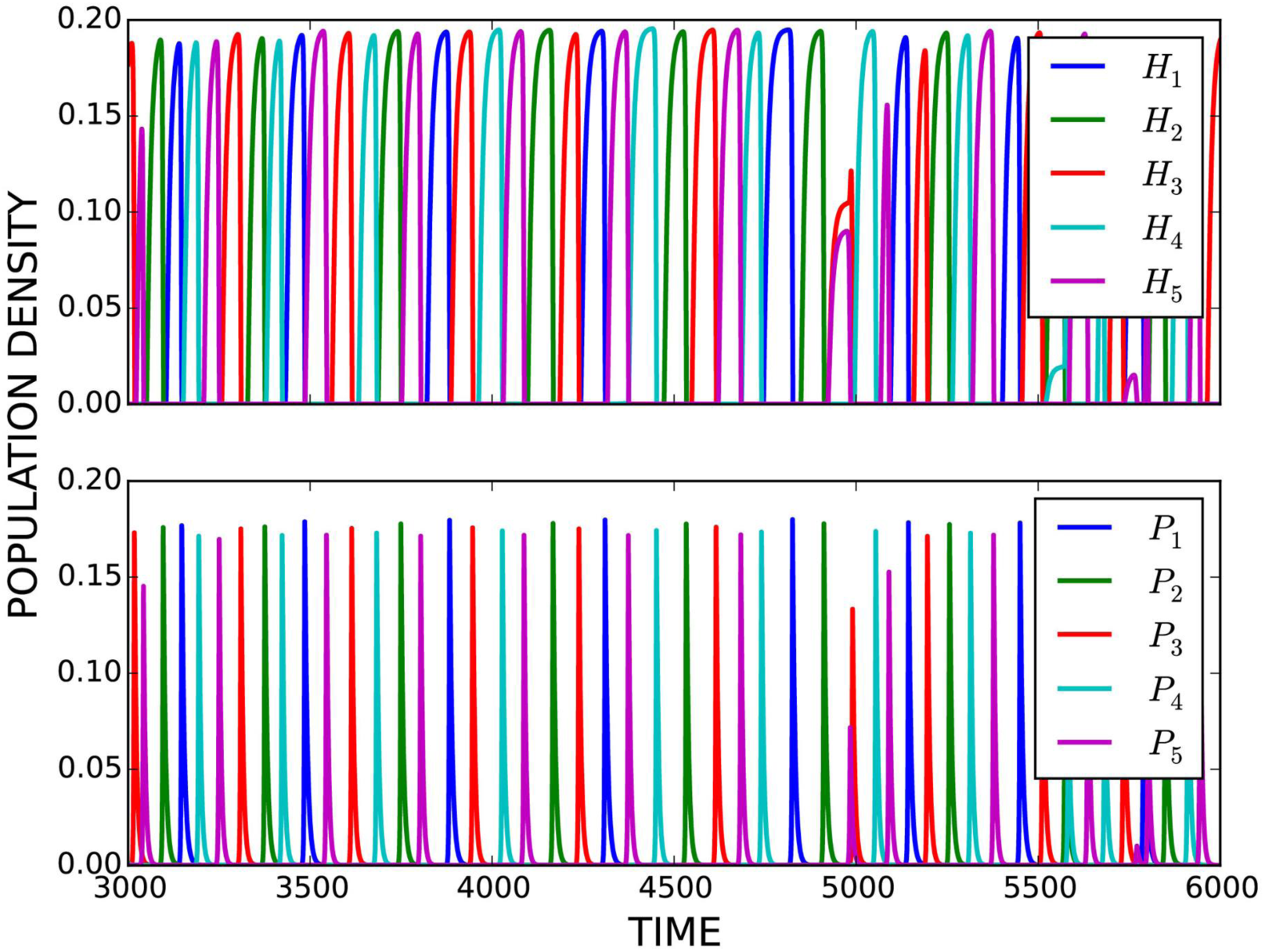
(**e**) Red Queen dynamics, a special type of oscillatory dynamics. The Red Queen dynamics is described by a time series of ever-changing dominant phenotype frequencies characterized by out-of-phase cycles and identical amplitude [Rabajante et al. 2015; Rabajante et al. 2016]. The cycles are pathogen-induced, that is, a common host type with high frequency is infected by pathogens resulting in the eventual decrease in frequency. As the frequency of the common host type decreases, a rare host type is given the opportunity to increase its frequency and become the new dominant type resulting in winnerless cycles. The winnerless cycles also happen in the pathogen types because pathogens tracktheir specific hosts.

In our simulation, fluctuating negative frequency-dependent selection, specifically the Red Queen dynamics, happens in the model where each host phenotype has an associated specific pathogen (Fig. 4e). This high pathogen specificity characterizes a host-pathogen interaction with tight matching of host and pathogen traits, as in the matching-alleles model [Weitz et al. 2012; Rabajante et al. 2015]. In the Red Queen dynamics occurring in our model (Fig. 4e), the occurrence of cycles that are out-of-phase and have identical amplitude is consistent with the definition of the canonical Red Queen dynamics [Brockhurst et al. 2014; Rabajante et al. 2015; Rabajante et al. 2016]. The canonical Red Queen dynamics state that the average fitness of species stays the same (represented by identical amplitude) even though the species undergo continuous evolution (represented by out-of-phase cycles). The out-of-phase cycles imply that only one species type is dominant for a certain period of time and the rest are rare; but after some time, the dominant type is replaced by a rare one (cyclic phenotype-switching). Notice that the Red Queen dynamics do not only include cyclic phenotype-switching in hosts but also cyclic phenotype-switching in pathogens (Fig. 4e). Such Red Queen dynamics is one of the many manifestations of the Red Queen hypothesis [Avrani et al. 2012; Brockhurst et al. 2014]. There are several empirical evidences based on experiments and fossil records supporting the Red Queen hypothesis [Brockhurst et al. 2014; Raberg et al. 2014; Voje et al. 2015]. Investigating cyclic coevolutionary dynamics could also help in designing strategies to manage drug resistance (for example, see Fig. 17 in [Mira et al.2015]).

In the point-of-view of cell and developmental biology, oscillations like the Red Queen dynamics may represent excessive temporal plasticity of cells. This excessive plasticity is one of the characteristics of a mutator (epimutator) phenotype that may lead to phenotypic diversity or to tumor heterogeneity in cancer [Shackleton et al. 2009; Marusyk & Polyak 2010; Loeb 2011; Peltomaki 2012; Swanton 2012]. Oscillations that characterize a mutator phenotype occur due to aberration in the gene regulatory network [Loeb 2011; Rabajante & Babierra 2015]. Here we have shown that pathogens (e.g., oncovirus) can also cause the occurrence of mutator phenotype [Liao 2006], especially when there is winnerless coevolution between host immune system and the pathogens (Fig. 4e). Abnormal dampening of these oscillations could lead to irreversible gene misexpression. In population biology point-of-view, the Red Queen dynamics show pathogen-mediated temporal diversity in host and pathogen species. However, the never-ending cycles in the Red Queen dynamics may not proceed indefinitely in ecological systems. This is because host-pathogen interactions are finite activities [Avrani et al. 2012; Brockhurst et al. 2014; Voje et al. 2015]. Sooner or later, some host and parasite types may escape the arms race competition. Several species may likewise become losers in the arms race competition (e.g., species extinction) because they were not able to cope up with the coevolutionary process.

### Epigenetic inheritance

One of the important components for bringing the outcomes of epigenetic landscape to the level of fitness landscape is epigenetic memory, which is characterized by epigenetic marks. Epigenetic memory is required in non-genetic inheritance of phenotypes from cell to cell or from organism to organism (transgenerational) [Pal & Miklos 1999; Bond & Finnegan 2007; Pilu 2011; Geoghegan & Spencer 2012; DUrso & Brickner 2014; English et al. 2015]. Much of the epigenetic memories are reset in the germline but some are conserved and imparted to offspring [Rakyan et al. 2001; Molinier et al. 2006; Bonduriansky & Day 2009; Robertson & Richards 2015]. Transgenerational inheritance happen in different timescales [Rando & Verstrepen 2007; Rabajante & Babierra 2015]. In some cases, epigenetic changes could lead to rapid phenotype variation in populations but epigenetic marks could also decay in few generations [Rando & Verstrepen 2007; Geoghegan & Spencer 2012].

In our decision-switch model, self-stimulation (positive feedback; Fig. 2) and multistability support epigenetic memory [Cinquin & Demongeot 2005; Kaufmann et al. 2007; Lim & van Oudenaarden 2007; Cheng et al. 2008; Smits et al. 2008; Hughes 2012]. Self-stimulation allows the expression of a phenotype to be fitter through time. In a multistable system, each phenotype attractor has their own basin of attraction. The probability of a phenotype being heritable is higher if (i) the phenotypes basin of attraction is large, that is, it is stable for a wide range of initial conditions, and (ii) the phenotype attractor is structurally stable, that is, it is stable for a wide range of parameter values. If the initial condition and parameter values are reset during meiosis, epigenetic marks can escape resetting if the new initial condition and parameter values still fall within the stability range of the phenotype attractor. Large basin of attraction and structural stability also provide robustness against environmental noise. Nevertheless, further studies are needed to map the extent of opportunities and limitations of epigenetic inheritance, especially on how it affects the genetics and evolution of populations [Crisp et al. 2016].

### Future prospects

Empirical investigations on the interface between pathogens, epigenetics and evolution are in progress [van Nhieu & Arbibe 2009; Cosseau et al. 2010; Thomas et al. 2011; Gomez-Diaz et al. 2012; Brockhurst & Koskella 2013]. Diversity of perspectives is useful but a unified perspective involving different fields, such as epigenetics, parasitology, pathology, epidemiology and evolutionary biology, will help better understand and solve problems about diseases [Gomez-Diaz et al. 2012; Noble 2015; Skinner 2015]. Our theoretical predictions could aid ongoing and future studies on identifying the non-genetic effect and mechanisms of diseases from the cellular to population level. Bacteria-phage interaction in marine ecosystems, plant-pathogen interaction in agricultural systems, drug dynamics in synthetic/artificial organisms, and epigenetic-based diseases in insects are some of the plausible subjects for experimental validation [Takken & Rep 2010; Koskella & Brockhurst 2014; Williams 2013; Gijzen et al. 2014; Mukherjee et al. 2015; Cini et al. 2015; Trozzet & Carbonell 2015]. In this paper, we have employed a minimal model to study the expression of phenotypes in cells and populations. Other scenarios can be explored further, such as the dynamics along the continuum between matching-alleles and gene-for-gene systems [Agrawal & Lively 2002; Weitz et al. 2012; Rabajante et al. 2015], the effect of the number of hosts and pathogens [Woolhouse et al. 2002; Rabajante et al. 2015], the effect of different infection networks (e.g., random, modular, nested) [Weitz et al. 2012], and the interplay among many genetic, epigenetic and environmental factors [Price et al. 2003; Mostowy & Engelstadter 2011; Soen 2014]. The epigenetic factors could include in detail the different mechanisms that alter gene expression, such as DNA methylation, chromatin remodeling, histone modifications and RNA interference [Rakyan et al. 2001; Bonduriansky & Day 2009; Rohlf et al. 2012; Duncan et al. 2014; Kilvitis et al. 2014; Brazel & Vernimmen 2016].

## Methods

### Definition of variables and parameters

Suppose we have *m* varieties of host phenotypes and *n* types of pathogens/parasites. There are two state variables in the model, *H*_*i*_ and *P*_*j*_. We can compute for *F(H_i_)* that denotes the population frequency of host phenotype *i*, where *F(H*_*i*_*)=H*_*i*_/*N*_*H*_, *N*_*H*_=Σ_*k*_*H*_*k*_ (*k=1,2,…,m*). *F(P_j_)* denotes the population frequency of pathogen strain *j*, where *F(P*_*j*_*)=P*_*j*_/*N*_*P*_, *N*_*P*_=Σ_*l*_*P*_*l*_ (*l=1,2,…,n*). Population frequency may also be interpreted as the probability that a phenotype will be expressed. For the definition of parameters, see Table 1. All state variables and parameters are non-negative.

**Table 1.** Definition of parameters.

| Parameter | Definition |
| --- | --- |
| $r_i$ | coefficient associated with the growth of host type $i$, or the efficiency in expressing phenotype $i$ |
| $K_i$ | factor influencing the carrying capacity allocated to host type $i$ |
| $\rho_i$ | death rate of host type $i$ not due to pathogens, or the decay rate of the expression of phenotype $i$ |
| $c_i$ | exponent affecting the curve of self-stimulation of host phenotype $i$; hyperbolic if $c_i=1$, sigmoidal if $c_i>1$. For simplicity, assume $c_i$ is integer-valued. |
| $c_{ik}$ | exponent affecting the nonlinear repression behavior of host phenotype $k \neq i$ against host phenotype $i$. For simplicity, we can assume $c_{ik}=c_i$ for all $i,k$. |
| $\theta_i$ | non-zero threshold constant; if all $H_{k \neq i}=0$, then $\frac{r_i H_i^{c_i}}{\theta_i + H_i^{c_i}} K_i = \frac{r_i K_i}{2}$ when $H_i = \theta_i^{1/c_i}$ |
| $\gamma_{ik}$ | interaction coefficient associated with the repression of host phenotype $i$ by host phenotype $k \neq i$ |
| $g_i$ | basal or constitutive growth rate (sometimes called the 'leak' term), i.e., the expression of host phenotype $i$ still increases at this rate at $H_i=0$ |
| $\alpha_{ij}$ | strength of pathogenicity or infectivity of pathogen strain $j$ against host type $i$ |
| $\varphi_{ij}$ | host $i$-to-pathogen $j$ conversion rate that defines the growth rate of pathogen population |
| $d_j$ | death rate of pathogen strain $j$ |
| $q_i$ | exponent defining the shape of the functional response curve. For simplicity, we can assume all host-pathogen relationships have the same functional response type. |
| $\phi_i$ | non-zero threshold constant of the functional response curve |
| $\beta_{ikj}$ | coefficient representing the energy allotment of pathogen strain $j$ to different host types; if $\beta_{ikj}=0$, $i \neq k$, then the population of host type $k$ is negligible in altering the functional response curve for infecting host type $i$ |

### Cellular decision-making model

The following epigenetic model (Eq. 1) describes the qualitative dynamics of multistable decision switches in cell-fate determination, based on the gene regulatory network in Fig. 2. Each node in Fig. 2 represents a set of gene regulatory factors involved in expressing a specific phenotype. The nodes have mutual repression because it is presumed that a mature specialized cell expresses a unique phenotype and inhibits the expression of the other phenotypes. For further details about this model and definition of variables and parameters in epigenetics point-of-view, refer to [Cinquin & Demongeot 2005; Macarthur et al. 2008; Andrecut 2011; Rabajante & Babierra 2015; Rabajante & Talaue 2015] (note that there are some differences in the symbols used).

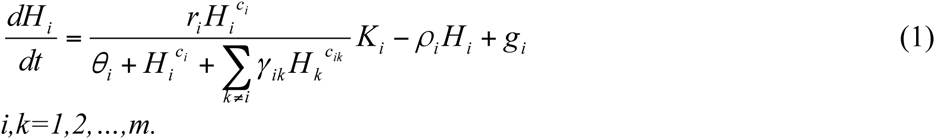

### Frequency-dependent selection

With proper definition of variables and reparametrization, we assume similar decision-making mechanism (Eq. 1) happens during evolutionary selection of a phenotype. If a phenotype is advantageous, a population of species may select this phenotype resulting in higher phenotypic fitness (characterized by a higher phenotype frequency) compared to the other phenotypes. This model is similar to the Lotka-Volterra competition model but involving a non-polynomial function that describes multistability and self-stimulation (auto-activation/auto-catalysis) [Rabajante & Talaue 2015]. Multistability and self-stimulation supports the preservation of epigenetic memory.

The battle for phenotype dominance is akin to inter-host competition. An increase in inter-host competition 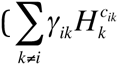 in the denominator of equation (1)) reduces the net effect of the host growth rate coefficient *r*_*i*_. The value 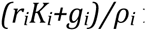 is an upper bound of the equilibrium value of *H*_*i*_. The parameter *K*_*i*_ affects the upper bound, which we can interpret as a factor influencing the carrying capacity allocated to each host type. Here we assume the allocation is fixed. The total carrying capacity is 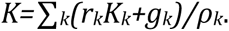

### Model of host-pathogen interaction

To model host-pathogen interaction, we couple equation 1 with pathogen dynamics. We assume that the pathogen dynamics follow a parasitism model, as follows [Smout et al. 2010; Jover et al. 2013; Rabajante et al. 2015; Rabajante et al. 2016]:

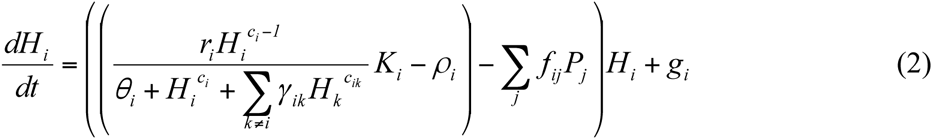

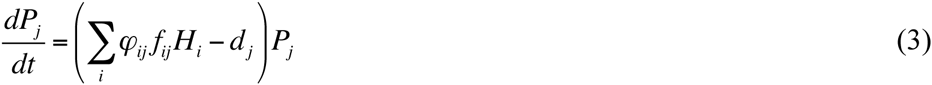

where

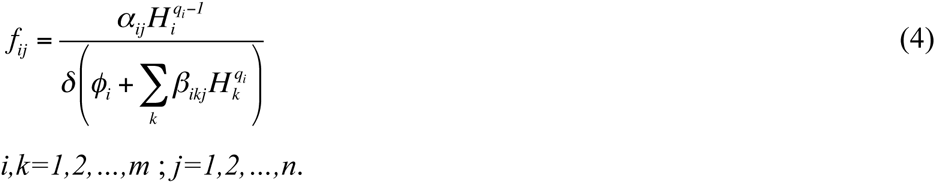

The population frequency of host phenotype is decreased by parasitism quantified by the functional response *ƒ*_*ij*_*H*_*i*_. The growth of pathogens depends on host utilization (numerical response 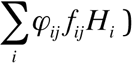) with death rate *d*_*j*_. Equation (4) defines the pathogen functional response. If *q_i_ =1* and then the functional 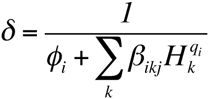 response curve is of type-I (linear). If *q_i_=1* and *δ=1* then the functional response curve is of type-II (hyperbolic). If *q_i_=2* and *δ=1* then the functional response curve is of type-III (sigmoidal).

In our simulations, we are primarily concerned with the dynamics of the epigenetic landscape. We assume a fixed set of genotypes (fixed structure of gene regulatory network). The predicted phenotype variation due to epigenetic factors under the influence of parasitism could be amplified if modifications in genetic factors are incorporated in our model.

## Acknowledgment

We would like to thank Juancho A. Collera of UP Baguio and Editha C. Jose of UPLB for discussion about the model and its analysis.

## Author contributions

MJVC ran the simulations and produced the results. JFR conceived the study and built the model. JMT created the figures. JFR, ALB, MJVC and JMT wrote the manuscript.

## Supplementary material

Parameter values used in the simulations

